# Natural variations of adult neurogenesis and anxiety predict hierarchical status of inbred mice

**DOI:** 10.1101/2024.09.21.614232

**Authors:** Fabio Grieco, Atik Balla, Thomas Larrieu, Nicolas Toni

**Affiliations:** Center for Psychiatric Neuroscience, Department of Psychiatry, Lausanne University Hospital, University of Lausanne, Prilly, Switzerland

## Abstract

Hierarchy provides a survival advantage to social animals in challenging circumstances. In mice, social dominance is associated with trait anxiety and reduced stress resilience which are regulated by adult hippocampal neurogenesis. Here, we tested whether adult hippocampal neurogenesis may regulate social dominance behavior. We observed that future dominant individuals exhibited higher trait anxiety and lower levels of hippocampal neurogenesis prior to social hierarchy formation, suggesting that baseline neurogenesis might predict individual social status among a group. This phenotype persisted after social hierarchy was stable. Experimentally reducing neurogenesis prior to the stabilization of social hierarchy in group-housed males increased the probability of mice to become dominant and increased anxiety. Finally, when innate dominance was assessed in socially isolated and anxiety-matched animals, mice with impaired neurogenesis displayed a dominant status toward strangers. Together, these results indicate that adult neurogenesis predicts and regulates hierarchical and situational dominance behavior along with anxiety-related behavior. These results provide a framework to study the mechanisms underlying social hierarchy and the dysregulation of dominance behavior in psychiatric diseases related to anxiety.

## Introduction

The hierarchical structure within animal societies arises from the necessity to address environmental challenges crucial for the group’s survival. In such circumstances, the formation of a hierarchy is as an effective mechanism to manage fluctuations in resource availability, mitigating conflicts, conserving energy, and fostering stability, thereby providing a strong survival advantage to the group. In humans, problems with dominance and social hierarchy are found across a broad range of psychopathologies: Externalizing disorders such as mania-proneness, and narcissistic traits are related to heightened dominance behaviors, whereas anxiety and depression are related to subordination and submissiveness^1, 2^. While high status in human and non-human primates can reduce stress when the social hierarchy is stable, instability can exacerbate stress responses in individuals with higher status^3,4^. Indeed, individuals who occupy dominant positions are often subjected to higher expectations, and increased responsibilities, all of which contribute to elevated stress levels, and the continuous effort to assert and defend their status in the face of potential challenges can be inherently stressful and anxiety-provoking^5^. In mice, hierarchical dominance is associated with anxiety-like behavior, with dominant mice displaying greater anxiety-related behavior than subordinates, leading to depressive-like symptoms^6–8^. However, the mechanisms underlying the interplay between mood-related behavior and social dominance remain elusive.

Alteration in adult hippocampal neurogenesis is one of the major features of anxiety disorder and depression^9^. Adult hippocampal neurogenesis (AHN) results in the continuous formation of neurons throughout life, that integrate into the hippocampal network^10^. These newly formed neurons play crucial roles in memory mechanisms and mood regulation, evidenced by the fact that inhibition of adult neurogenesis reduced memory capabilities along with increased stress vulnerability to depression^11–14^. Interestingly, the impairment of adult neurogenesis results in increases in anxiety-related behavior^15^. Considering that dominant individuals display emotional alterations under baseline conditions, we hypothesized that this behavioral phenotype can be associated with alterations in hippocampal adult neurogenesis and that innate inter-individual differences in adult neurogenesis may underlie differences in trait anxiety and dominance behavior. We tested this hypothesis using a combination between an observational study and two interventional studies aimed at inhibiting adult neurogenesis.

## Results

### Future dominant mice display lower levels of baseline adult hippocampal neurogenesis and heightened trait anxiety than subordinates

We first assessed the relationship between social hierarchy, adult hippocampal neurogenesis and anxiety. To do so, 6-week-old male mice were individually housed and tested for trait anxiety using elevated plus maze (EPM), light-dark (LDT) and open field (OFT) tests, followed by 5-Iodo-2’-Deoxyuridine (IdU) injection to label proliferating cells, enabling the evaluation of baseline neurogenesis (Fig. 1A). The mice were then housed in groups of 4 individuals per cage, and anxiety levels were reassessed using the same tests after one week of cohabitation, a period corresponding to the active establishment of social hierarchy. The mice were left undisturbed for an additional four weeks (i.e., for a total of 5 weeks of cohabitation), which is sufficient to enable the establishment of a stable hierarchy^7^, and anxiety was tested again. 5-Chloro-2′-deoxyuridine (CldU) was administered 24h after anxiety assessment to assess the potential influence of social hierarchy on adult neurogenesis. Finally, individual social ranks were assessed using a social confrontation tube test (SCTT, which reflects cage social hierarchy^16^) in a round robin design for 2 weeks. In an independent cohort, we observed that the hierarchical rank in the SCTT was stabilized after 3 days (Fig. EV 1A-C). Importantly, the social rank assessed in the tube test was neither linked to the time spent in the tube during the 2-day habituation phase (Fig. EV 1E) nor to the body weight before or after the tube test (Fig. EV 1F-H). We observed that before cohabitation, mice that later became dominant exhibited higher anxiety score than subordinate mice (Fig. 1B). This was reflected by a reduction in the time spent exploring the open arms of an EPM, the lit compartment in a LDT and the center during an OFT (Fig. EV 2A-D). Notably, comparable findings were obtained after 1 and 5 weeks of cohabitation (Fig. 1C-D, Fig. EV. E-J), suggesting that the anxiety profile observed in high-ranking mice is not solely attributable to the active agonistic behavior during the first week of hierarchy establishment but instead results from a pre-existing, individual anxiety trait.

**Figure 1:**
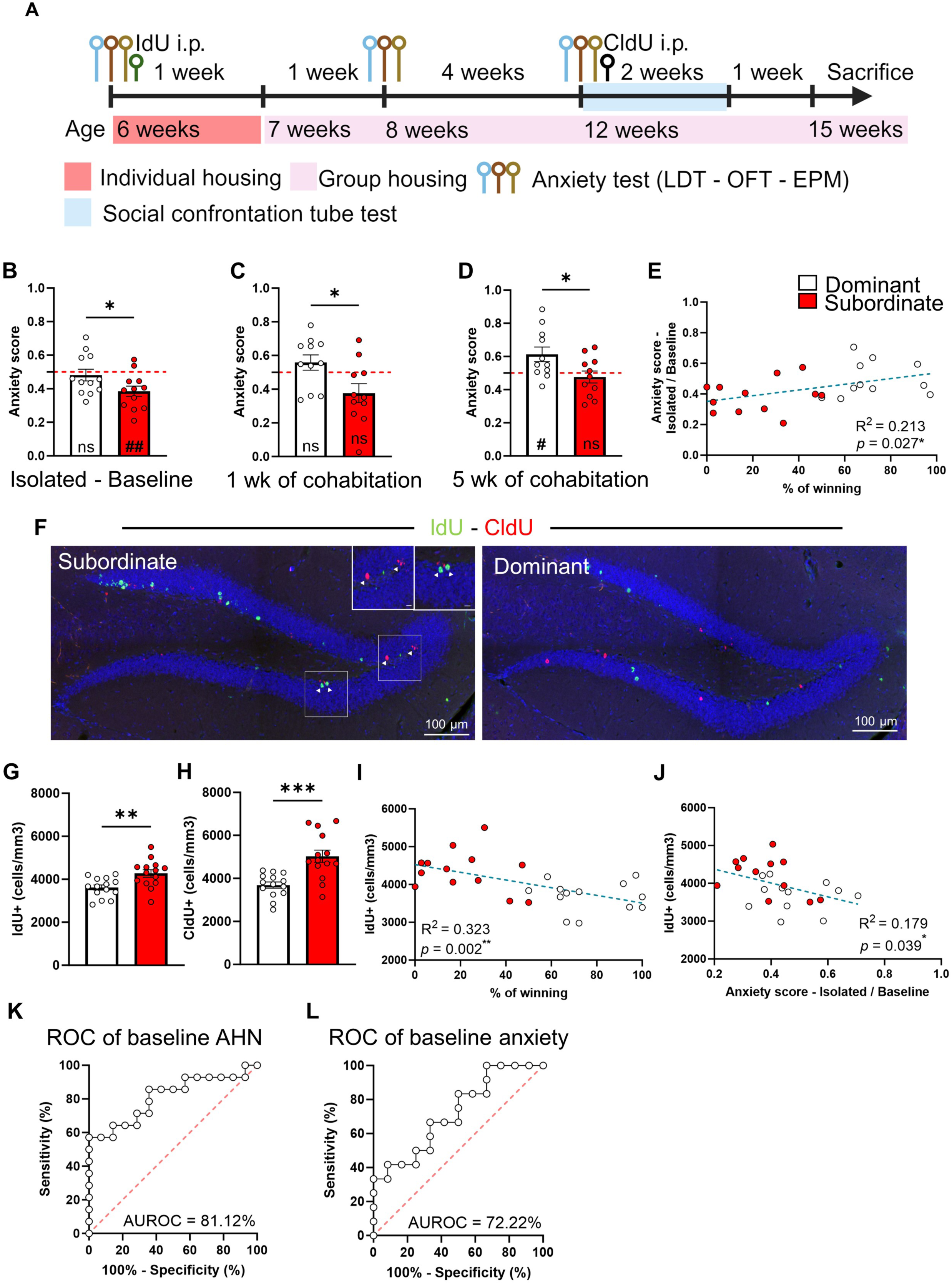
Dominant mice display persistent anxiety traits and decreased AHN. **A.** Schematic illustration of the experimental design for the evaluation of anxiety, adult hippocampal neurogenesis and social rank. **B-C-D.** Anxiety scores in dominant and subordinate mice during isolation (**C**, baseline) after 1 (**C**) and 5 (**D**) weeks of cohabitation (anxiety score derived from normalization of time spent in the dark chamber during LDT, time spent into the closed arm in the EPM, and time spent in thigmotaxis in the OFT. (**B**, t22=2.07, p = 0.50, unpaired t test, two-tailed, n = 12 per group; t11=0.53 dominants, p = 0.602, one sample t test, n = 12; t11=3.71 subordinates, p = 0.003, one sample t test, n = 12. **C**, t20=2.52, p = 0.020, unpaired t test, two-tailed, n = 11 per group; t10=1.29 dominants, p = 0.225, one sample t test, n = 11; t10=2.20 subordinates, p = 0.052, one sample t test, n = 11. **D**, t20=2.38, p = 0.027, unpaired t test, two-tailed, n = 11 per group; t10=2.54 dominants, p = 0.02, one sample t test, n = 11; t10=0.65 subordinates, p = 0.525, one sample t test, n = 11). **E.** Correlation between baseline anxiety score and percentage of winning during SCTT (R^2^ = 0.2133, p = 0.027, simple linear regression). **F.** Confocal maximal projections of the dentate gyrus, immunostained for IdU (green), CldU (red) and DAPI (blue) in subordinate and dominant animals. **G-H.** Quantification of CldU-(**G**) and IdU-positive cells (**H**) in the dentate gyrus (**G**, t26=4.15, p = <0.001, Welch’s test, two-tailed, n = 14 per group; **H**, t26=3.10, p = 0.005, unpaired t test, two-tailed, n = 14 per group). **I-J.** Correlations between IdU counts and percentage of winning (**I**) or baseline anxiety score (**J**) (**I**, R^2^ = 0.323, p = 0.002, simple linear regression; **J**, R^2^ = 0.179, p = 0.039, simple linear regression). **K-L.** Receiver Operating Characteristic (ROC) curve for the discrimination of dominant individuals by baseline AHN (**K**) and baseline anxiety (**L**). Histograms show average ± SEM, * p<0.05, ** p<0.01, *** p<0.001, ns = not significant. Comparison between the group mean and the hypothetical value of 0.5, to assess anxiety withing each group are shown within each histogram bar. One-sample t-test. #: p<0.05; ##: p<0.01, ns = not significant. Inset scale bars: 10µm.

When comparing anxiety between the three time points (baseline and after 1 and 5 weeks of cohabitation), anxiety levels increased over time, as revealed by a one-sample t-test analysis comparison with the 0.5 value of anxiety score. This increased anxious phenotype appearing during the cohabitation period suggests that the ongoing maintenance of a social status, even within stable social hierarchies, imposes continuous stress that may contribute to elevated anxiety over time in both dominant and subordinate individuals. Finally, the proportion of winning confrontations in the SCCTT linearly correlated with anxiety-related behavior at baseline (Fig. 1E) and after 1 week of cohabitation, where higher anxiety was associated with higher social status but not after social hierarchy stabilization (Fig. EV 3A, B).

Adult hippocampal neurogenesis plays a role in mood regulation^11–14, 17, 18^ and anxiety^15^. To assess whether hierarchy was associated with adult hippocampal neurogenesis, we analyzed CldU- and IdU-immunolabeled cells. Dominant individuals exhibited significantly fewer IdU-labeled and CldU-labeled cells than subordinates (Fig. 1G-H). Furthermore, the number of IdU cells inversely correlated with both the proportion of wins (Fig. 1I) and anxiety at baseline (Fig. 1J) but not after 1 week and 5 weeks of cohabitation. Similarly, the number of CldU-labeled cells inversely correlated with the proportion of confrontations won (Fig. EV 3C-E). Thus, hierarchical rank is associated with variations in adult neurogenesis and anxiety, that are present before and are maintained after the establishment of hierarchy.

These observations suggests that baseline neurogenesis and trait anxiety may predict future social status within a group. To test his possibility, we used a Receiver Operating Characteristic (ROC) curve analysis. Baseline hippocampal neurogenesis accurately distinguished dominant from subordinate animals with 81% accuracy (Fig. 1K), whereas baseline anxiety-like behavior showed a 72% accuracy in segregating dominant from subordinate individuals (Fig. 1L). Together, these results suggest that adult neurogenesis and, to a lesser degree anxiety, are pre-existing traits that play a determinant role in hierarchy establishment.

### Inhibiting adult neurogenesis increases social dominance behavior and anxiety

Social dominance is expressed by either hierarchical (confrontation between cage mates) or situational (confrontation between strangers) dominance behavior ^19, 20^. To assess the role of adult neurogenesis in hierarchical dominance behavior, mice were housed in groups of 4 individuals. Starting after 2 days of cohabitation, while the social hierarchy was still being established, two of the four mice were treated either with temozolomide (TMZ), an antimitotic drug that impairs adult neurogenesis^21, 22^ or vehicle (saline, NaCl 0.9%) for the first 3 days of each week for four consecutive weeks, followed by 2 weeks of rest. To assess cell proliferation, mice were injected with BrdU after the first week of TMZ administration. After this, mice were tested in the SCTT using a round robin design, followed by anxiety assessments (Fig. 2A). Consistent with previous studies^21, 22^, we observed that TMZ treatment reduced the number of BrdU-positive cells (Fig. 2B, C). Furthermore, mice subjected to TMZ treatment demonstrated a significant increase in the probability of acquiring a dominant status in the SCTT (Fig. 2D). We also observed a significant increase in the number of wins in TMZ-treated compared to NaCl-treated mice (Fig. 2E, F). Finally, we found that TMZ-treated mice showed increased anxiety score as compared to NaCl-treated mice (Fig. 2G, Fig. EV 4). Overall, these results suggest that adult neurogenesis regulates anxiety and social hierarchy in the home cage.

**Figure 2:**
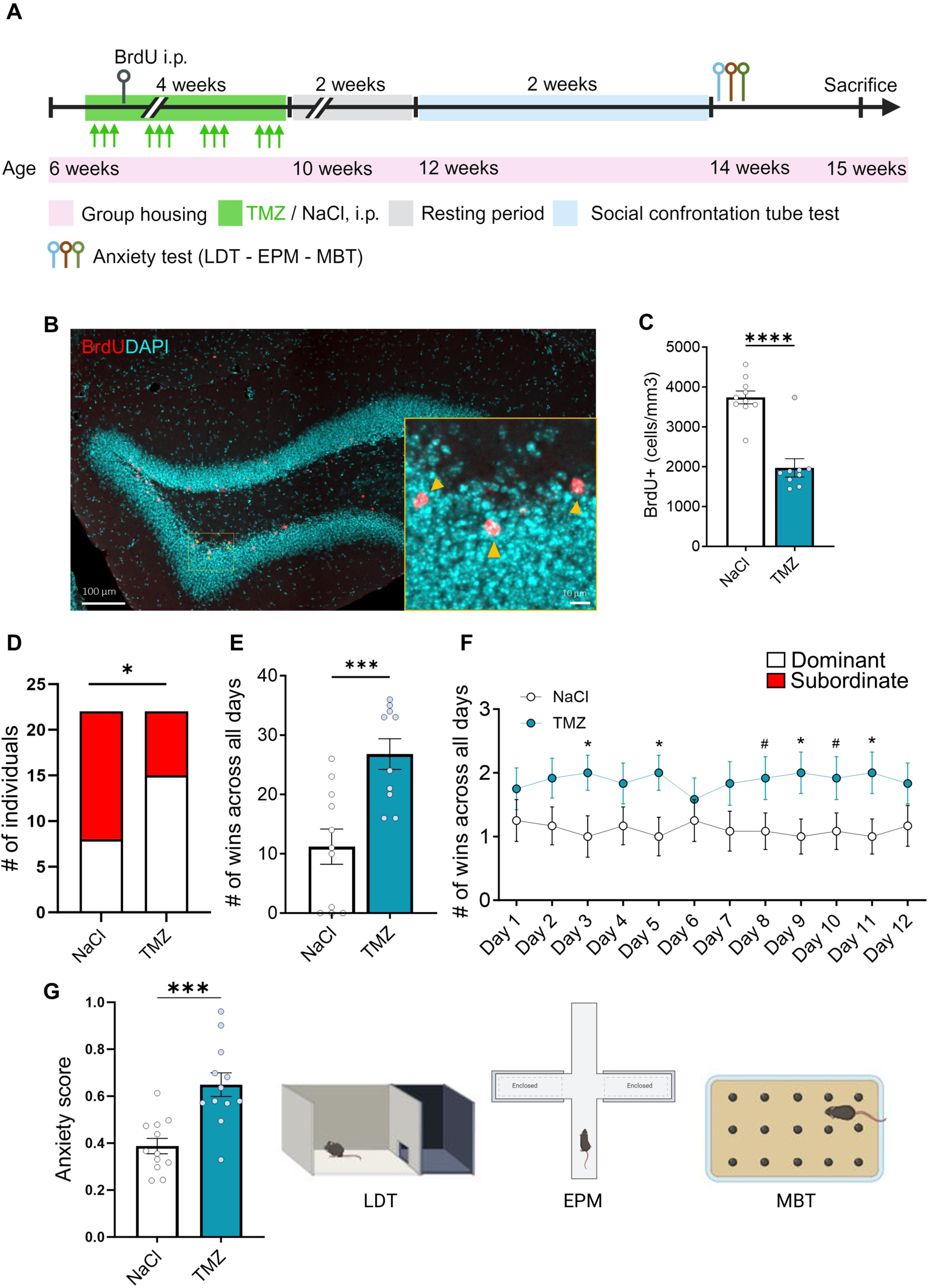
Adult neurogenesis depletion increases hierarchical dominance. **A.** Experimental design for TMZ administration, anxiety assessment and SCTT of group-housed mice. **B.** Representative confocal microscopy image of the dentate gyrus immunostained for BrdU (red cells highlighted with yellow arrows) and DAPI (blue). **C.** BrdU-positive cell density in the dentate gyrus (t17 = 6.42, p = <0.0001, unpaired t test, two-tailed n = 10-9 per group). **D.** Contingency table for the effect of TMZ treatment on dominance behavior (Chi-square, two-sided, p = 0.0346). **E.** Histogram representing the number of wins in SCTT across all days (t19 = 3.92, p = 0.0009, unpaired t test, two-tailed, n = 11-10 per group). **F.** SCTT outcomes day by day in number of wins (interaction effect: F11,242 = 1.53, p = 0.104, time effect: F11,242 = 0.12, p = 0.99, mouse effect: F1,22 = 3.58, p = 0.071, subject effect: F22,242 = 69.84, p = < 0.0001; two-way ANOVA; 12-12 per group; in Day 3: t22 = 2.34, p = 0.028, unpaired t test, two-tailed, n = 12-12 per group; in Day 5: t22 = 2.44, p = 0.023, unpaired t test, two-tailed, n = 12-12 per group; in Day 8: t22 = 1.88, p = 0.073, unpaired t test, two-tailed, n = 12-12 per group; in Day 9: t22 = 2.34, p = 0.028, unpaired t test, two-tailed, n = 12-12 per group; in day 10, t22 = 1.88, p = 0.073, unpaired t test, two-tailed, n = 12-12 per group; in Day 11: t22 = 1.88, p = 0.073, unpaired t test, two-tailed, n = 12-12 per group). **G.** Histogram of the anxiety score of TMZ and NaCl-treated mice (anxiety score derived from normalization of time spent in the dark chamber during LDT, time spent into the closed arm in the EPM, and number of marbles buried in MBT: t22 = 4.36, p = <0.001, unpaired t test, two-tailed, n = 12-12 per group). Histograms show average ± SEM, * p<0.05, *** p<0.0002, **** p<0.0001.

Wild mice live in territories occupied by a single adult male, several females and their progeny, and therefore, male mice establish dominance upon their first encounter^23^. As such, situational dominance seems ethologically more relevant than hierarchical dominance observed in male mice sharing a cage. To assess the role of adult neurogenesis in situational dominance, we tested the effect of TMZ in a second cohort of male mice which were singly housed, to prevent the formation of social hierarchy. One week after arrival, mice were weighed and assessed for trait anxiety using an elevated plus maze, a light dark test and an open field test. The combined anxiety score (data not shown) was used to assign mice to either the TMZ or the vehicle group to match weight and anxiety between groups. After this, mice were treated with either TMZ or saline for 4 weeks, followed by 4 weeks of resting period. After this, their trait anxiety was assessed using an open-field, an elevated plus maze and a light-dark tests. TMZ-treated mice were then paired with saline-treated mice of similar anxiety traits and body weight and confronted in the SCTT for 5 days, twice a day (morning and afternoon, Fig. 3A). We found that across all sessions, TMZ-treated animals exhibited an increased probability of becoming dominant as compared to vehicle-treated animals (Fig. 3B). In addition, when considering the competition across the 5 testing days, TMZ-treated mice showed an increased number of total wins and wins per day (Fig. 3C, D). Although the anxiety score was similar between groups (Fig. 3E, Fig. EV 5), the anxiety score of the vehicle group was higher than in group-housed mice (Fig. 2G), likely reflecting the anxiogenic effect of long-term social isolation^24, 25^. Thus, inhibiting adult neurogenesis increased social hierarchy and dominance behavior in a situational context.

**Figure 3:**
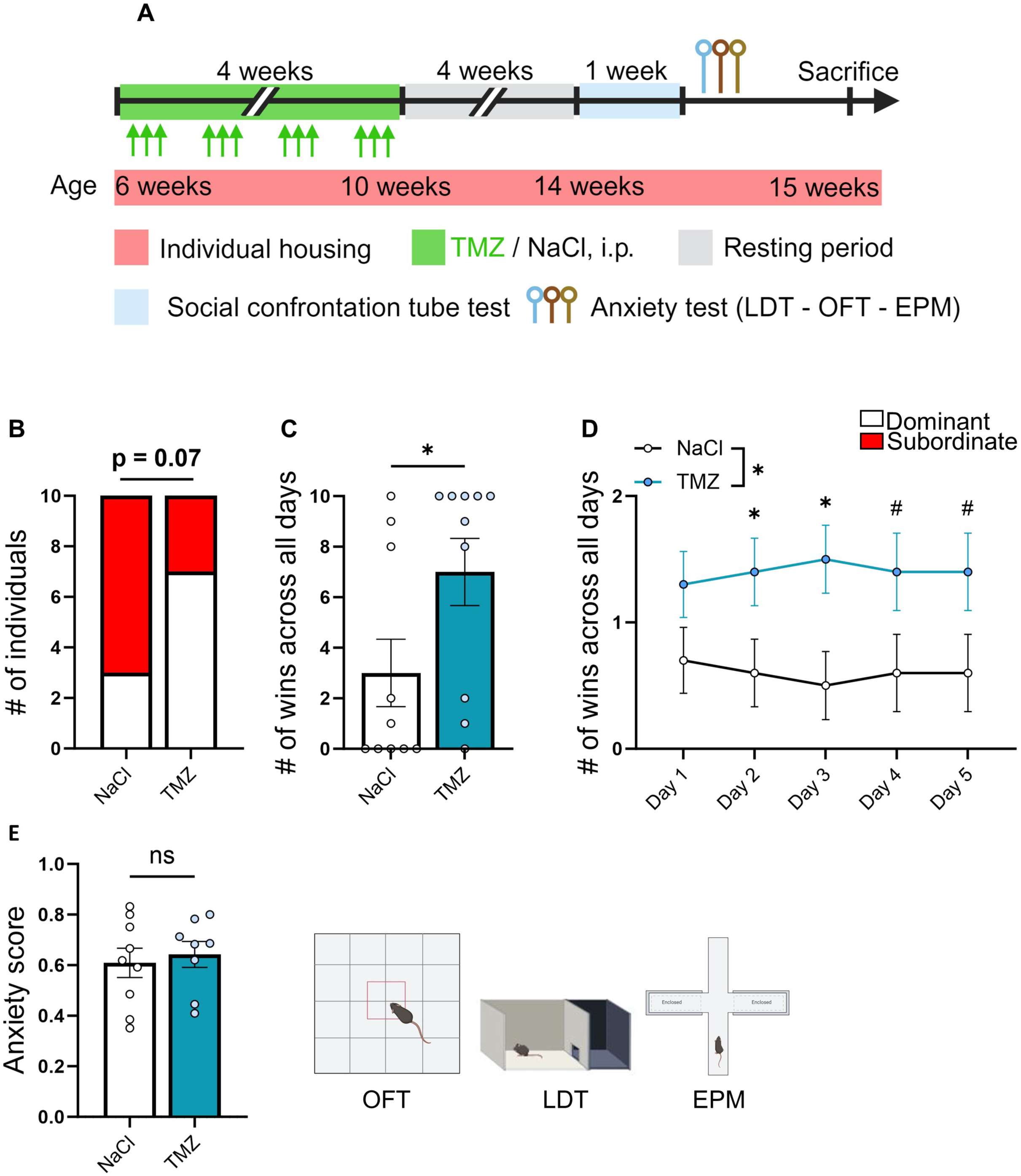
Adult neurogenesis depletion increases situational dominance. **A.** Experimental design for TMZ administration, anxiety assessment and SCTT in individually housed animals. **B.** Contingency table for correlation between tube test ranks and solution administrated. (Chi-square, two-sided, p = 0.073). **C.** Histogram representing the number of wins in SCTT across all days (t18 = 2.12, p = 0.0480, unpaired t test, two-tailed, n = 10 per group). **D.** SCTT outcomes day by day in number of wins. (interaction effect: F4,72 = 0.94, p = 0.441; time effect: F2,111 = 0.00, p = > 0.999; mouse effect: F1,18 = 4.50, p = 0.048; subject effect: F18,72 = 33.68, p = <0.0001; two-way ANOVA; n = 10-10 per group; in Day 2: t18 = 2.12, p = 0.048, unpaired t test, two-tailed, n = 10 per group; in Day 3: t18 = 2.63, p = 0.016, unpaired t test, two-tailed, n = 10 per group; in Day 4: t18 = 1.85, p = 0.080, unpaired t test, two-tailed, n = 10 per group; in Day 5: t18 = 1.85, p = 0.080, unpaired t test, two-tailed, n = 10 per group). **E.** Histograms of the anxiety score derived from OFT, LDT and EPM tests. Histograms show average ± SEM, * p<0.05. ns=not significant.

## Discussion

In this study, we examined the role of adult hippocampal neurogenesis in shaping social dominance behavior and trait anxiety. First, we found that before group formation, individuals with higher trait anxiety and lower levels of hippocampal neurogenesis became dominant when groups were formed. After housing the mice in groups, dominant animals continued to show heightened anxiety levels and less neurogenesis than subordinates both during and after hierarchy establishment, indicating that hierarchy formation did not modify these pre-existing traits. We next assessed the role of adult neurogenesis in the regulation of social dominance behavior. We found that reducing neurogenesis increased anxiety under baseline conditions, as well as dominant behavior hierarchical conditions, i.e. between recent cage mates that are establishing hierarchy, indicating that adult neurogenesis plays a role in the maintenance of hierarchy. Furthermore, inhibiting adult neurogenesis in mice that had never met, also increased dominance in confrontation tests, supporting the view that neurogenesis plays a role in the establishment of hierarchies. Interestingly, TMZ-treated mice displayed dominant behavior towards anxiety-matched vehicle-treated mice, suggesting that the effect of neurogenesis on hierarchy is not mediated by anxiety. Together, these results indicate that adult neurogenesis regulates trait anxiety as well as dominance behavior, even in absence of prior social experience and suggest that anxiety is not induced by dominant behavior but is rather a pre-existing trait. In addition to providing a mechanistic link between adult neurogenesis, anxiety and dominance behavior, these results offer a framework to study social behavior in the context of psychiatric diseases.

The relationship between adult neurogenesis and social dominance is subject to strong inter-species, sex, and methodological differences^26–33^. For example, previous studies in rats using the visible burrow system, an environment designed to stimulate dominance hierarchy formation within a complex spatial and social context, have found an increased number of new-born neurons in dominant rats compared to their subordinate counterparts^28, 29^. In contrast, a study of naked mole rats in an intact colony, showed that dominant breeders exhibited significantly reduced DCX immunoreactivity in the dentate gyrus of the hippocampus compared to subordinates^34^. In mice however, studies where social hierarchy was determined either by video recordings of agonistic behavior in the home cage^31^ or on social confrontation tube test^26^, revealed no discernible difference in adult neurogenesis between dominant and subordinate individuals.

Here, our findings revealed that individuals who later became dominant within a social group exhibited notably higher levels of trait anxiety along with reduced hippocampal neurogenesis even before the formation of social structures. This pre-existing neurobiological profile suggests that baseline neurogenesis may serve as a predictor of an individual’s future social status within a group. The stability of this profile over time suggests that individuals with lower baseline neurogenesis and higher anxiety may be neurobiologically predisposed to assert dominant positions within social hierarchies. However, while our findings clearly establish the role of adult neurogenesis in the initial formation of social hierarchies, its involvement in the maintenance and regulation of dominance behavior within an already-established hierarchical group remains to be explored.

Furthermore, it remains unclear how decreased adult neurogenesis induces dominance. The effect of TMZ on dominance may be mediated by a reduction of neurogenesis in the olfactory bulb or in the hippocampus or both. Indeed, neurogenesis reduction experiments have shown that both olfactory and hippocampal neurogenesis play a role in social recognition (the discrimination between a familiar and a new mouse) and social memory^30, 35–37^. The vomeronasal system has also been shown to play a role in dominance behavior^26, 38^. Thus, a reduction of social memory may stimulate the appearance of novel agonistic behavior, leading to the establishment of a novel hierarchy. However, in the present study, TMZ also increased dominance behavior in non-familiar mice, therefore in absence of social memory. Although the local reduction of adult neurogenesis may reveal the respective contributions of olfactory bulb and hippocampal neurogenesis to dominance behavior, the results presented here suggest that the effect of adult neurogenesis on dominance is mediated by mechanisms independent of social recognition.

The effect of reduced neurogenesis on dominance may be mediated by anxiety. The relationship between dominance and anxiety behavior in rodents is complex and multi-factorial and may depend on individual differences in adult neurogenesis, contextual factors, adaptive responses, or environmental constraints. Moreover, the relationship between anxiety and dominance behavior may be bidirectional. Indeed, the continuous pressure to maintain high social status is associated with increased stress and HPA axis activation upon challenges^3^. Inversely, trait anxiety may lead to dominance as an adaptive strategy to a challenging environment. For example, in the presence of an anxiogenic context, such as an aquatic barrier that limits access to food, a group of rats develop social differentiation in which dominant, non-diving rats steal food from diver, subordinate rats^39^. Furthermore, the direct regulation of dominance by anxiety has been tested by the administration of anxiolytics showing that the acquisition of a subordinate position correlated with a decrease in anxiety levels^40^. Although our observational study support this hypothesis, the finding that TMZ increases dominance in animals that were anxiety-matched suggests that while anxiety and dominance behavior can be influenced by overlapping neural circuits, anxiety by itself may not be totally causative of dominance behavior, but rather a consequence of an underlying phenotype. Thus, based on this finding, it is likely that the effect of adult neurogenesis on dominance is independent from anxiety.

Several brain regions are known to be involved in the perception and learning of social dominance, such as the nucleus accumbens^7, 41^, the ventral tegmental area^42^ and the prefrontal cortex^19, 43^. In particular, the medial prefrontal cortex (mPFC) seems to play a predominant role in social hierarchy behavior: the winning history of individual mice remodels thalamic connections to the mPFC, leading to long-term modifications of the dominance status^44^. Furthermore, the excitatory synaptic efficacy in the mPFC is higher in dominant mice than in subordinates, and the bidirectional manipulation of synaptic efficacy in the mPFC alters social dominance^43, 44^. In this context, it is important to keep in mind that in our experiments, TMZ was injected systemically and may therefore affect other mechanisms than neurogenesis, which may also participate to the dominance behavior. However, adult neurogenesis is a particularly drastic mechanism of plasticity that enables a rapid adaptation to novel living conditions. Interestingly, the ventral hippocampus projects to the mPFC and the activity of these projections is involved in social memory^45^ as well as in the response to anxiogenic experience^46^. Owing to their specific connectivity, immature adult-born hippocampal neurons decrease the activity of mature neurons from the ventral dentate gyrus in a stressful environment^14^. Thus, one may speculate that reduced adult neurogenesis may promote dominance behavior by increasing the output from the ventral hippocampus to the mPFC.

Due to their crucial role in individual survival, allocation of territory, access to reproduction and food resources, dominance behaviors stand out as pivotal constraints influencing the social organization and structure of a group. The reduced neurogenesis in dominant mice may confer behavioral advantage in their natural habitat: In the wild, *Mus musculus* live in demes formed by one dominant adult male that defends the deme’s territory, several females, and their offspring. As soon as the male offsprings reach sexual maturity, they will either remain in the deme under the dominance of the adult male until they inherit the territory or leave the original deme to form their own^47, 48^. In addition, in group foraging scenarios, individuals vary in their commitment to seeking food and their utilization of food discovered by others. In the framework of these naturalistic considerations, one can speculate that individuals with higher neurogenesis will display lower anxiety and higher spatial memory and are therefore more likely to explore new territories to form their own deme or to identify new food source. In contrast, individuals with lower neurogenesis and higher trait anxiety may remain in and defend the deme’s territory and may choose to steal food from others, a strategy for which dominance behavior provides an advantage. In this context, pre-existing inter-individual differences in hippocampal neurogenesis may be one of the mechanisms underlying inter-individual differences in anxiety^15^, stress response and dominance behavior^7, 8, 49–51^. Interestingly, adult neurogenesis is strongly influenced by numerous environmental conditions such as physical activity, aging, nutrition, diseases, anxiety and stress^17, 18, 52–54^. Thus, despite the genetic homogeneity of inbred mice, the life history of an individual leaves a trace in the hippocampal connectivity which, in turn, influences the mouse behavior, including dominance behavior.

In humans, dominance behavior plays an important role in the formation of hierarchies. A disruption of dominance behavior is associated with several psychiatric disorders such as conduct disorders, antisocial personality disorder, and certain forms of depression and anxiety^1, 2^. The identification of adult neurogenesis as an important regulator of dominance behavior not only enhances our understanding of the regulation of dominance behavior, but also provides a novel venue for intervention for impaired dominance behavior in the context of psychiatric disorders in humans.

## Acknowledgements

The authors wish to thank the confocal imaging facility of the University of Lausanne (CIF) for access and training to confocal microscopy. This project was funded by the Swiss National Science Foundation (Grant Number 310030_201015).

## Author contributions

FG, TL, and NT designed research. FG, TL, and AB performed research. FG, TL, AB, and NT analyzed the data. FG, TL, and NT wrote the paper. TL and NT supervised the research. NT obtained funding for the project.

## Materials and methods

### Animals

All experiments were performed according to the ARRIVE guidelines^55^ and with the approval of the local Veterinary Authorities (Vaud, Switzerland, authorization number: VD3728) and carried out in accordance with the European Communities Council Directive of 24 November 1986 (86/609EEC). All experiments were performed on C57Bl/6J male mice obtained from Janvier Laboratories (France). After arrival at 6 weeks of age (to avoid excessive aggressivity seen in older adult mice), animals were housed four per cage (social hierarchy experiments) or in individual cages (situational dominance) and allowed to acclimate to the vivarium for one week before the start of experiments. During that time, all animals were subsequently handled for 1 min. per day for a minimum of 3 days. Animals were weighted upon arrival as well as weekly to monitor health. Mice were maintained under standard housing conditions on corncob litter in a temperature-(23 ± 1°C) and humidity-(40%) controlled animal room with a 12h. light/dark cycle (8h00–20h00), with unlimited access to food and water. Retired CD1 breeders used as unfamiliar social targets in the social interaction test were obtained from Charles River laboratories. All tests were conducted during the light period.

### TMZ and BrdU/CldU/IdU administration

To assess the basal level of adult hippocampal neurogenesis IdU (57.5 mg/kg; 10mg/ml) was dissolved in a 0.2N NaOH/saline solution and injected i.p., three times the same day, starting from 11 am with 2 hours interval between each injection. Then, mice were injected with a CldU solution (42.5mg/kg; 10mg/ml i.p.), three times the same day with 2 hours interval between each injection. To reduce neurogenesis, mice were treated with TMZ as previously described^21, 56^. An intraperitoneal (i.p.) injection of either TMZ (25 mg/kg; 2.5 mg/ml in 0.9% NaCl) or saline (0.9% NaCl) was given for the first three days of the week for a total of 4 weeks, followed by another 4 weeks of resting period between the last injection and the following behavioral test. Moreover, in preparation for the assessment of cell survival via immunostaining, the mice were injected with BrdU after the first cycle of TMZ. Each animal was injected with BrdU (100 mg/kg; 10 ml/kg i.p.) three times the same day, starting from 11 am with 2 hours interval between each injection.

### Behavioral tests

#### Elevated plus maze test

The test was conducted as previously described^7^. Briefly, the animals were placed in a maze made from black PVC with a white floor. The EPM consists of 4 arms (30 x 5 cm) arranged in a plus shape, with walls on two opposing arms, while the other two arms are exposed to the height of the apparatus (65 cm). The arms converge at the center, forming a small, squared platform (5 x 5 cm). Throughout the tests, the light conditions were maintained stable. The open arms and the center of the maze had a luminous flux of 12-15 lux, while the closed arms had reduced light intensity with 5 lux. The animals were gently introduced into the maze, facing the wall at the end of the closed arms, and were allowed to freely move within the maze for a duration of 5 minutes. Video recordings of the mice’s behavior were captured from above the arena, and tracking analyses were performed using the ANY-maze software to determine the time spent in both the open and closed arms.

#### Open-field test / Novel object tests

The OFT was conducted as previously described^7^ in a rectangular arena (50 × 50 × 40 cm3) illuminated with dimmed light (30 lux). Mice were introduced near the wall of the arena and allowed to explore for 10 min. Analyses were performed using ANY-maze tracking software by drawing a virtual zone (15 × 15 cm2) in the center of the arena defined as the anxiogenic area. Several parameters were analyzed, including the total distance travelled and the time spent in the different zones. At the end of the 10 minutes, an object (4 x 4 x 15 cm) was placed in the center of the apparatus and mice could freely explore the novel object during 5 minutes in the same conditions as previously.

#### Light - Dark test

As previously described^57^, the apparatus utilized for the LDT consisted of a white wooden box with two compartments. One was a square compartment without a lid, serving as the light side (40 x 40 cm), while the other was a smaller rectangular compartment with a lid, creating the dark side (20 x 40 cm). These two compartments were connected by a 5 x 5 cm door, and the entire apparatus had a height of 30 cm. The center of the lit compartment maintained a stable luminosity of 400 lux, while the dark compartment remained without any light source. Mice were introduced into the apparatus in the light side, facing the door, and allowed to explore for a duration of 5 minutes. The mice’s movements were tracked and recorded using the ANY-maze software. In this test, anxiety-like behavior was evaluated based on the time spent in the dark compartment.

#### Marble burying test

The experiment was conducted following the methodology described^58, 59^. The apparatus used for the test consisted of an open, transparent plastic box measuring 40 x 25 x 20 cm, filled with bedding that had a depth of 6 cm. The test involved placing 20 dark marbles, each 16 mm in diameter, spaced evenly in a 4 x 5 grid on the surface of the bedding. Mice were placed in the middle of the box and were free to explore for 20 minutes under 300 lux. Upon completion of the experiment, the mice were removed from the box, and the number of buried marbles was visually assessed. A marble was considered “buried” when more than two-thirds of its surface was covered by the bedding. The evaluation of an elevated number of buried marbles is indicative of an anxious profile in the mice.

#### Social confrontation tube test

The social confrontation tube test (SCTT) was conducted on mice cohabitating for a duration of 5 weeks. Each mouse underwent training to cross a clear Plexiglas tube (diameter: 3cm; length: 30cm) five times from each end over two consecutive days. The tube’s diameter allowed unidirectional movement for adult mice. During the habituation phase, retreat or cessation of movement prompted gentle encouragement by touching the tail with a plastic stick. Between trials, the tube was cleaned with a 70% ethanol solution to eliminate smell, urine, or feces. Following the two-day habituation, social ranks were assessed over 9 consecutive days. Before the confrontation phase, each mouse was retrained to cross the tube from each end. Using a round-robin design, pairwise confrontations were conducted within social groups, resulting in six trials per cage of four mice. Two mice were simultaneously guided by the tail at the opposite tube ends until they reached the middle of the tube, and the time spent in the tube was recorded until one mouse compelled its cage mate to retreat. The mouse that retreated was identified as the ‘loser’ for that specific trial. Subsequently, for each trial, the same mouse alternated being placed in the tube from each end. Social dominance was quantified by calculating the percentage of winning time, and mice were ranked from 1 to 4, with Ranks 1 and 2 signifying the most dominant mice and Ranks 3 and 4 indicating the most subordinate mice.

The SCTT was also used to evaluate situational social dominance (as opposed to hierarchical status). Mice from each treatment group (TMZ or NaCl) were paired with mice from the other group, attempting to match body weight and trait anxiety. The couples were tested twice a day, in the morning and afternoon. To prevent potential confounding effects and biases, we took precautions to avoid pairing innate dominant saline mice with innate subordinate TMZ mice. Following an initial random pairing, the outcome yielded dominant mice from both the TMZ and saline groups. In a subsequent pairing, we specifically matched dominant TMZ mice with dominant saline mice that never met before with similar body weight and trait anxiety. The second round of confrontation revealed the social dominance of mice based on the number of wins across the 5 days. Mice with a percentage of victories higher than 50% were considered dominant. The Plexiglas tube was cleaned between each trial with 70% ethanol to remove odors, urine, and feces.

#### Anxiety score

The anxiety scores were a simplified version of principal component-based analyses^60, 61^ and encompassed several anxiety tests to obtain a general profile of anxiety, as previously described^6, 7, 62, 63^. The scores were derived from the normalization of the values for the combination of individual anxiety tests (time spent in the dark chamber during LDT, time spent into the closed arm in an EPM, time spent in thigmotaxis in an OFT and marbles buried). The normalization involved adjusting each animal’s value by subtracting the minimum value of the entire population and then dividing this result by the difference between the maximum and minimum values of the entire population: (x – min value) / (max value – min value). This method generates scores distributed on a scale from 0 to 1, with a score of 1 indicating high anxiety. The rationale for employing different combinations of tests was to increase the robustness of our findings by exploring whether TMZ would consistently influence anxiety-like behavior across a broader range of validated paradigms. While the open field, light-dark, and elevated plus maze tests are all well-established tools for assessing anxiety, we included additional measures, such as the marble burying test in the second experiment, to account for potential variations in anxiety responses that different tasks might detect. This approach also allowed us to confirm the reliability of TMZ’s effect by showing that any observed behavioral changes were not limited to a specific subset of tests but rather generalizable across multiple anxiety paradigms. Normalizing the results across three tests in each experiment ensures consistency in our scoring method, even as we explore diverse anxiety tests. The variation of the test combinations enhances the rigor of the study without compromising the validity of the anxiety score.

### Experimental designs

#### Experiment 1

We investigated the possible interplay between adult hippocampal neurogenesis, anxiety and dominance behavior. Six-week-old male mice were isolated after their arrival, after three days of acclimatization we tested their basal anxiety level using the EPM, the LDT and the OFT. After the last test, the mice were injected with 5-Iodo-2’-Deoxyuridine to evaluate baseline levels of adult hippocampal neurogenesis. After one week isolated, we formed anxiety matched group made by 4 animals per cage. We repeated the anxiety tests (EPM, LDT and OFT) one week after the formation of the cages to observe if the establishment of the social hierarchy was influencing their anxiety traits.

After a total of 5 weeks of cohabitation, when the social hierarchy is established and can be considered stable, we evaluated again their anxiety levels repeating the same battery of tests (EPM, LDT and OFT). At the end of the last test, mice were injected with 5-Chloro-2′-deoxyuridine. Together with the anxiety tests at different time points, the injections of thymidine analogues made possible the investigation of the link between basal levels of anxiety and adult hippocampal neurogenesis with the social hierarchy formation and establishment. After five weeks of cohabitation with their cage mates, mice underwent a SCTT to determine individual social rank within the cage for 3 weeks, in order to discriminate between dominants (rank 1 and 2) and subordinates (rank 3 and 4). The experimenter was blind to the identity of the mice during data collection and analyses. After the end of all behavioral tests, animals were sacrificed in the morning and their brains dissected out and prepared for histology.

#### Experiment 2

We investigated the role of adult neurogenesis in the establishment of social hierarchy in a group of four six-week-old male mice. After two days of cohabitation, TMZ was randomly administered to 2 mice from the cage while the other two received NaCl. One week following the initiation of TMZ treatment, mice received an intraperitoneal injection of BrdU to evaluate adult neurogenesis in the DG at the end of the experiment. After four weeks of TMZ treatment and an additional 2 weeks of resting period, the mice underwent SCTT to assess individual social rank. Four weeks after the completion of the TMZ treatment, mice underwent behavioral tests, including LDT, EPM and MBT to evaluate anxiety. The experimenter was blind to the identity of the mice during data collection and analyses. Twenty-four hours after the last behavioral test, the mice were sacrificed, and their brains were removed after perfusion and stored at −20 °C for further processing.

#### Experiment 3

We investigated the role of adult neurogenesis in innate social dominance behavior in a situational dominance context. Six-week-old male mice were individually housed upon arrival to prevent the development of a hierarchical structure and social memory. After the acclimatization week, the mice underwent four weeks of TMZ treatment, followed by a four-week resting period. Subsequently, animals were tested for dominance with the SCTT. The experimenter was blind to the identity of the mice during data analyses, but mouse identity was revealed during behavioral experiments, to enable the matching between TMZ- and vehicle-treated mice. After completion of all behavioral tests, mice were sacrificed, and their brains were removed after perfusion. The brain tissue was then stored at −20 °C for further processing.

### Histology

After all the behavioral experiments the mice were sacrificed with a lethal injection of pentobarbital (10 mL/kg, Sigma-Aldrich, Switzerland) and transcardially perfused with saline solution (NaCl 0.9%) followed by 4% paraformaldehyde (PFA) solution (Sigma-Aldrich, Switzerland). Brains were removed and postfixed with PFA 4% at 4°C overnight. Then brains were transferred in a 30% sucrose solution for 3 days before being frozen at −20°C until slicing. Coronal frozen sections of a thickness of 40 μm. were sliced with a microtome or cryostat (microtome, Leica Microsystems; cryostat, Leica CM3050S) to obtain hippocampal sections conserved in a cryoprotectant solution (30% glycerol, 30% ethylene glycol and 40% PBS 1M) at −20°C until immunofluorescence staining.

### Immunofluorescence staining

One out of 6 slices containing hippocampal tissue were chosen to cover the whole dentate gyrus and used for immunostaining. For detection of BrdU-, IdU- and CldU-positive cells, immunohistochemistry was performed using a formic acid pre-treatment (formamide 50%, 10% SSC 20X and 40% MilliQ water) at 60°C for 2h followed by DNA denaturation in 2 M HCl for 30 min at 37°C and rinsed in 0.1 M borate buffer pH 8.5 for 15 min and 6 times in PBS 0.1M for 10 minutes. The slices were incubated in blocking solution (0.3% TritonX-100 and 10% horse serum in PBS) at room temperature for 1 h and then incubated under agitation at 4° C overnight in a blocking solution containing the following primary antibodies: Rat anti-BrdU/CldU (Abcam, 1:1000, ab6326), and Mouse anti-IdU monoclonal antibody (Abcam, 1:500, ab181664). After being washed again in PBS, sections were incubated for 2 hours in either of the following secondary antibodies: Alexa Fluor 594 goat anti-rabbit (Invitrogen, 1:300, A11032), and Alexa Fluor 594 goat anti-rat (Invitrogen, 1:300, A11007). Sections were finally incubated for 10 minutes in 4′,6-diamidino-2-phenylindole (DAPI, 2 μg/ml in PBS) solution to mark the cell nuclei. Sections were mounted onto glass slides and cover-slipped using FluorSave (Millipore). Imaging was performed using a Nikon NI-E Spinning disk microscope. BrdU-, CldU- and IdU-positive cells were counted using the NIS elements Nikon software (ver 6.02) and automatized supervised counting. The experimenter was blind to the identity of the mice. The density of BrdU-, CldU- and IdU-positive cells was quantified by dividing the number of BrdU+ cells by the corresponding volume of DG expressed in mm^3^.

### Statistical analyses

Statistical analyses were carried out with Prism 9 (GraphPad Software, San Diego, CA 92108, USA), using an alpha level of 0.05. We eliminated outlier values that deviated more than 2 standard deviations from the average. All data are presented as mean ± SEM. Data were tested for normality using the Shapiro-Wilk-test. For normally distributed measures, we used an unpaired, two-tailed Student t-test to estimate differences between the two groups in trait anxiety (EPM, OFT, LDT, MBT), in body weight, in social dominance (SCTT) and in hippocampal neurogenesis (CldU-, IdU- and BrdU-immunolabeled cells). To evaluate the effect of TMZ on social hierarchy establishment and situational social dominance a Chi-square test has been performed. The social-confrontation tube test results were obtained by using either an unpaired, two-tailed Student t-test to estimate differences between the two groups or a two-way analysis of variance ANOVA with repeated measure, with treatment and days as fixed factors. Analyses were followed by a Bonferroni post-hoc test when it was appropriated. To correlate the cell survival with the anxiety scores and percentage of winning, a Pearson linear regression analysis was carried out.

### Data availability

All data presented in this manuscript has been deposited on Zenodo: 10.5281/zenodo.13821490

**Figure EV 1: Hierarchical Rank Using a Social Confrontation Tube Test. A.** Example of one cage representing the tube test ranks and winning times as a function of tube test trials. **B.** Average of the ranks for the nine cages over the 9-day test trials. **C.** Time spent in the tube as a function of the rank pairing. (F5,48 = 49.69, p < 0.001, one-way ANOVA, n = 9 per rank pairing). **D.** Percentage of winning times for each rank after the 9 days of social confrontations (F3,32 = 494.6, p < 0.0001, one-way ANOVA, n = 9 per rank pairing). **E.** Average of the time spent in the tube for the 2 days of habituation phase as a function of the final social rank (interaction: F3,64 = 0.41, p = 0.7442; Rank effect: F3,64 = 2.44, p = 0.0719; Side effect: F1,64 = 7.787, p = 0.0983; two-way ANOVA, n = 9 per group). **F-G.** Body weight of mice as a function of their social rank before (**F**) and after (**G**) the social confrontation tube test (before: t34 = 0.89, p = 0.376, unpaired t test R1-R2 vs R3-R4, two-tailed, n = 18-18 per group ; after: t34 = 0.30, p = 0.761, unpaired t test R1-R2 vs R3-R4, two-tailed, n = 18-18 per group). **H.** Body weight evolution in dominant and subordinate mice. Histograms show average ± SEM, * p<0.05, ** p<0.01, *** p<0.001, ns=not significant.

**Figure EV 2: Detailed anxiety values in dominant and subordinate individuals**. **A.** Schematic illustration of the tests used to assess anxiety. **B-E-H.** Percentage of time spent in the closed arm during the EPM at baseline (**B**), after 1 week of cohabitation (**E**) and after 5 weeks of cohabitation (**H**) (**B**, t20=1.89, p = 0.073, unpaired t test, two-tailed, n = 11 per group; **E**, t21=2.02, p = 0.056, unpaired t test, two-tailed, n = 12 per dominants, n = 11 per subordinates; **H**, t20=1.54, p = 0.138, two-tailed, n = 11 per group). **C-F-I.** Percentage of time spent in the dark chamber during the LDT at baseline (**C**), after 1 week of cohabitation (**F**) and after 5 weeks of cohabitation (**I**) (**C**, t20=2.29, p = 0.033, unpaired t test, two-tailed, n = 11 per group; **F**, t22=2.09, p = 0.048, unpaired t test, two-tailed, n = 12 per group; **I**, t20=2.164, p = 0.043, unpaired t test, two-tailed, n = 11 per group). **D-G-J.** Percentage of time spent in thigmotaxis during the OFT at baseline (**D**), after 1 week of cohabitation (**G**) and after 5 weeks of cohabitation (**J**) (**D**, t21=2.44, p = 0.023, unpaired t test, two-tailed, n = 12 per dominants, n = 11 per subordinates; **G**, t21=2.39, p = 0.026, unpaired t test, two-tailed, n = 11 per dominants, n = 12 per subordinates; **J**, t22=2.18, p = 0.040, unpaired t test, two-tailed, n =12 per group). * p<0.05, ns=not significant.

**Figure EV 3: Correlations between AHN, anxiety and social dominance. A-B.** Correlations between anxiety and percentage of winning during the SCTT after one week of cohabitation (**A**) and five weeks of cohabitation (**B**). (**A,** R^2^ = 0.257, p = 0.016, simple linear regression; **B**, R^2^ = 0.144, p = 0.090, simple linear regression). **C.** Correlation between the number of CldU-positive cells and percentage of winning during the SCTT (R^2^ = 0.332, p = 0.004, simple linear regression). **D-E.** Correlation between the number of IdU-positive cells and the anxiety level after one week (**D**) and five weeks of cohabitation (**E**) (**D**, R^2^ = 0.093, p = 0.166, simple linear regression; **E**, R^2^ = 0.074, p = 0.220, simple linear regression). **F.** ROC curve for the discrimination of dominant individuals by a composite score made by the normalized values of IdU-positive cells in the DG and baseline anxiety traits.

**Figure EV 4: Effect of TMZ on anxiety after group formation. A.** Graphic representation of the behavioral tests used to assess anxiety. **B-C-D.** Percentage of time spent in the closed arm during EPM (**B**), in the dark chamber in LDT (**C**) and of marbles buried in MBT (**D**) (**B**, t21=3.55, p = 0.0019, unpaired t test, two-tailed, n = 11 and n = 12 per group; **C**, t18=2.87, p = 0.010, unpaired t test, two-tailed, n = 10 per group; **D**, t22=2.15, p = 0.042, unpaired t test, two-tailed, n = 12 per group). Histograms show average ± SEM, * p<0.05, ** p<0.01.

**Figure EV5: Effect of TMZ on anxiety before group formation. A.** Graphic representation of behavioral tests used to assess anxiety. **B-C-D.** Percentage of time spent in the closed arm during EPM (**B**), in the dark chamber in LDT (**C**) and in thygmotaxis during OFT (**D**) (**B**, t18=0.75, p = 0.458, unpaired t test, two-tailed, n = 10 per group; **C**, t18=0.36, p = 0.720, unpaired t test, two-tailed, n = 10 per group; **D**, t18=0.40, p = 0.690, unpaired t test, two-tailed, n = 10 per group). Histograms show average ± SEM, ns=not significant.

