## Supplementary figures and images for "Natural variations of adult neurogenesis and anxiety predict hierarchical status of inbred mice"

### Supplementary Figures

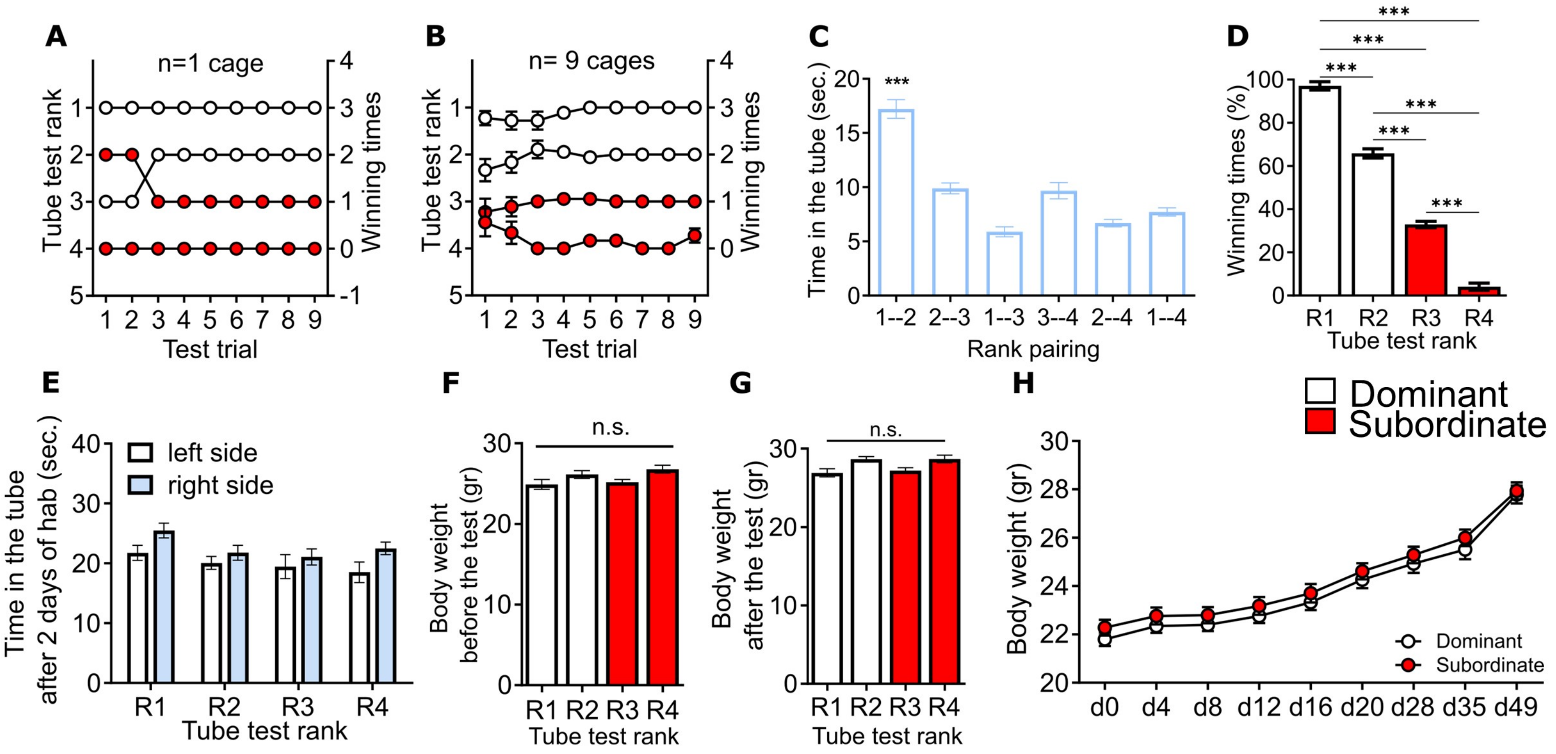

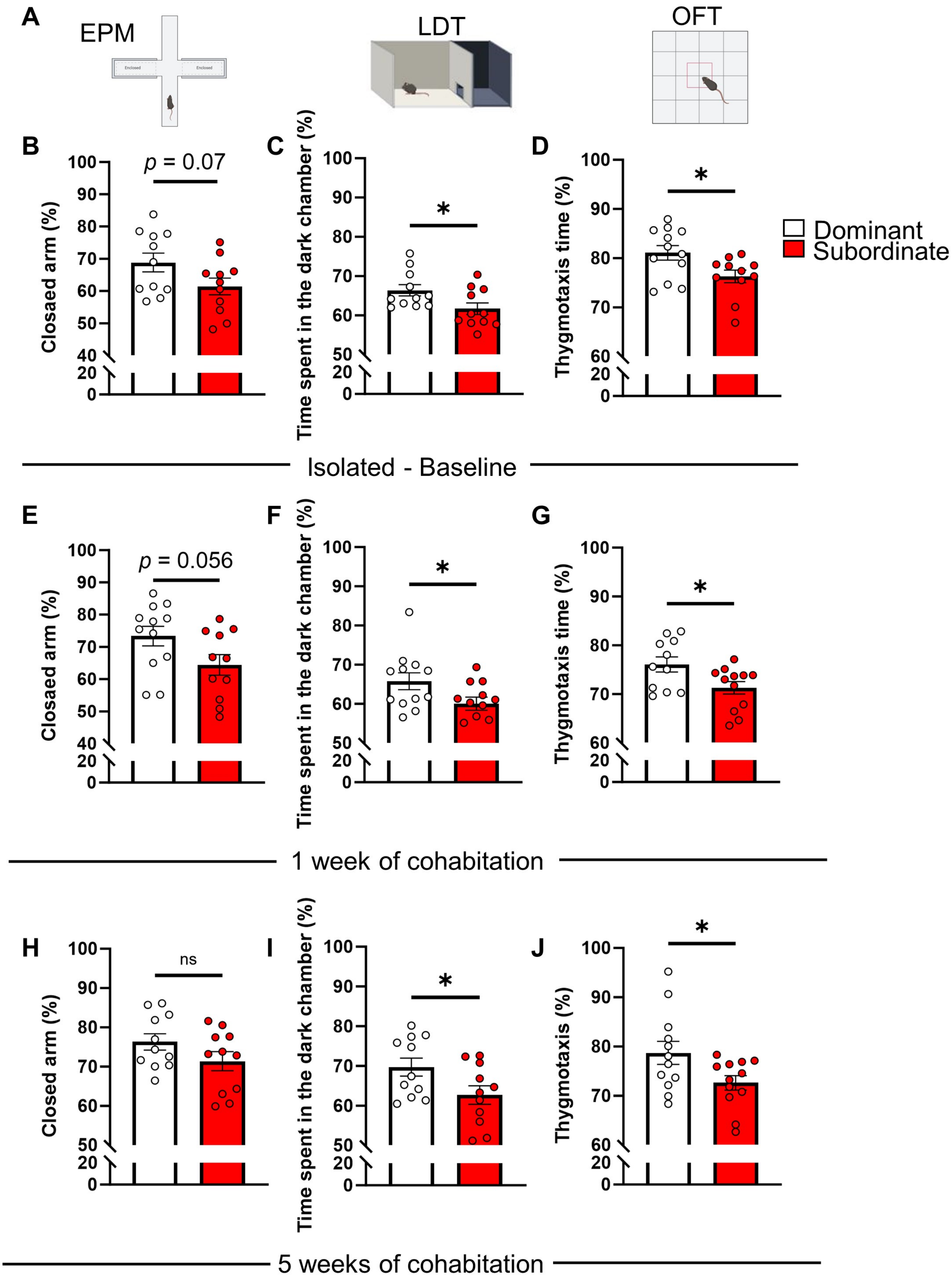

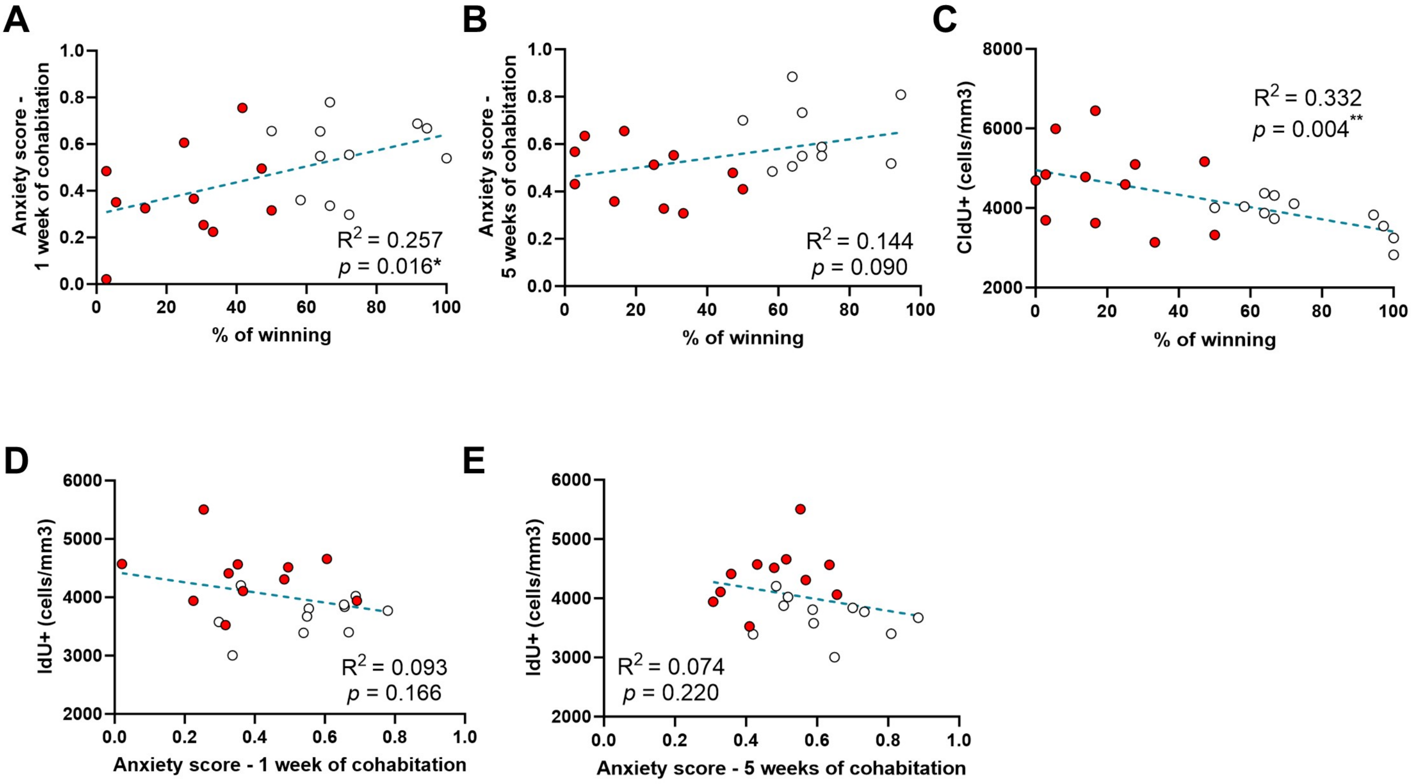

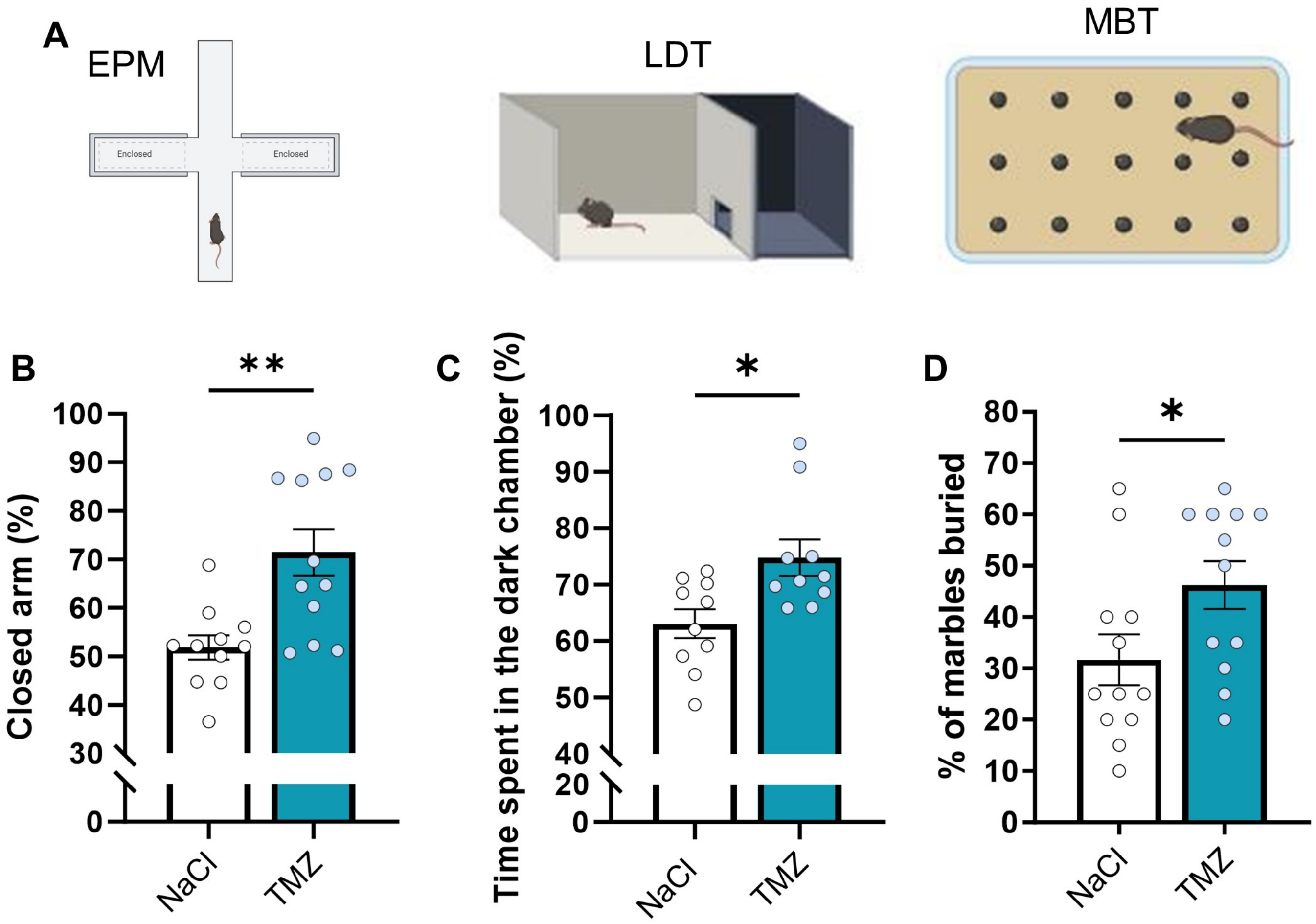

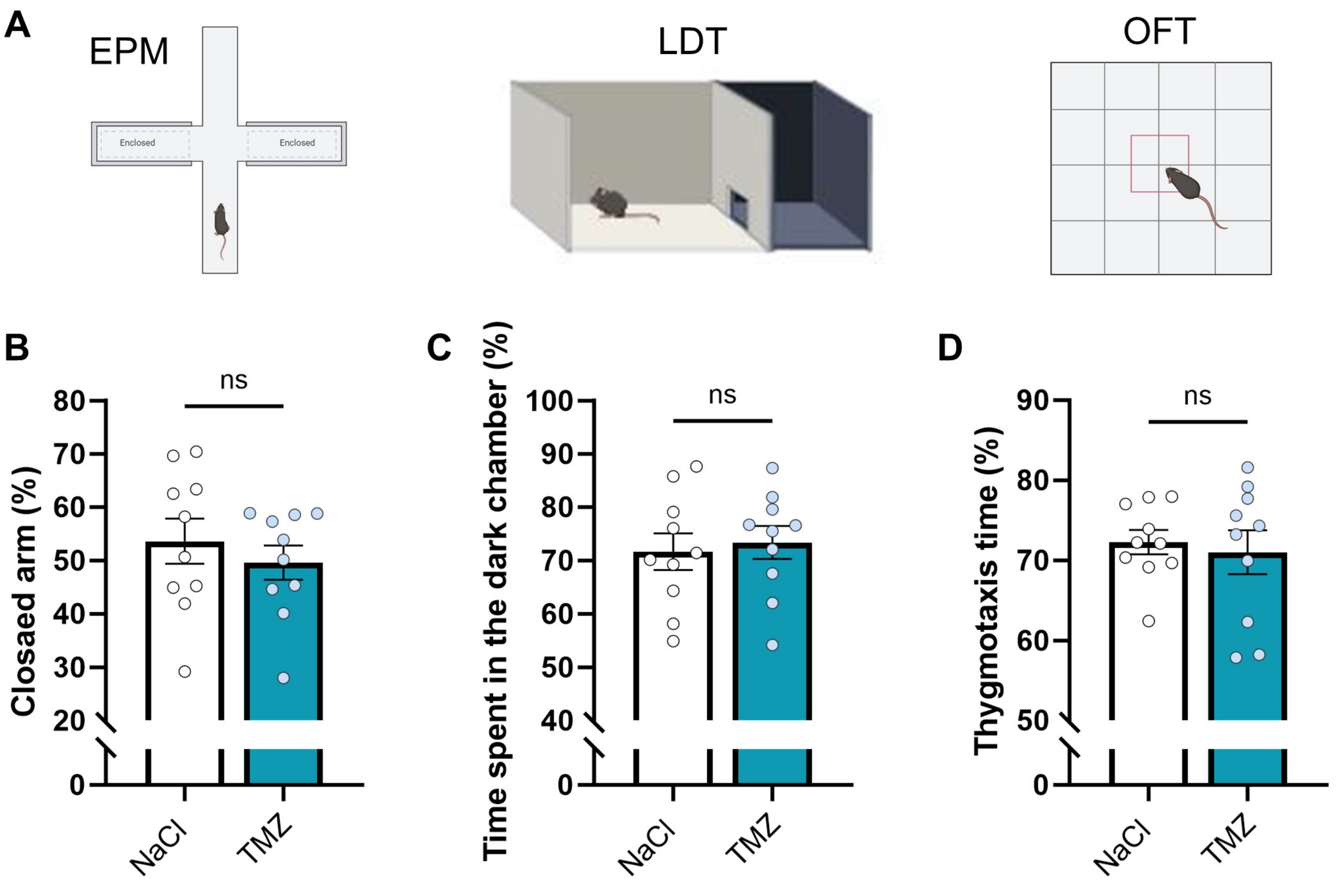
